# Glutamatergic heterogeneity in the neuropeptide projections from the lateral hypothalamus to the mouse olfactory bulb

**DOI:** 10.1101/2025.02.16.638511

**Authors:** Meizhu Qi, Julia Won, Catherine Rodriguez, Douglas A. Storace

**Author notes:** Corresponding Author: Douglas A. Storace, 107 Chieftan Way Florida State University Tallahassee, FL 32306. **Data Sharing and Data Availability:** Data will be made available upon request from the corresponding author. **Conflict of Interest:** The authors declare no financial or scientific conflicts of interest. **Ethics Approval Statement:** All authors understood the ethical principles that *The Journal of Comparative Neurology* operates under, and the work complied with the animal ethics checklist reported by (Grundy, 2015).

## Abstract

The direct pathway from the lateral hypothalamus to the mouse olfactory bulb (OB) includes neurons that express the neuropeptide orexin-A, and others that do not. The OB-projecting neurons that do not express orexin-A are present in an area of the lateral hypothalamus known to contain neurons that express the neuropeptide melanin-concentrating hormone (MCH). We used virally mediated anterograde tract tracing and immunohistochemistry for orexin-A and MCH to demonstrate that the OB is broadly innervated by axon projections from both populations of neurons. Orexin-A and MCH were expressed in each OB layer across its anterior to posterior axis. Both orexin-A and MCH neurons are genetically heterogeneous, with subsets that co-express an isoform of vesicular glutamate transporter (VGLUT). We used high-resolution confocal imaging to test whether the projections from orexin-A and MCH neurons to the OB reflect this glutamatergic heterogeneity. The majority (∼57%) of putative orexin-A axon terminals overlapped with VGLUT2, with smaller proportions that co-expressed VGLUT1, or that did not overlap with either VGLUT1 or VGLUT2. In contrast, only ∼26% of putative MCH axon terminals overlapped with VGLUT2, with the majority not overlapping with either VGLUT. Therefore, the projections from the lateral hypothalamus to the OB are genetically heterogeneous and include neurons that can release two different neuropeptides. The projections from both populations are themselves genetically heterogeneous with distinct ratios of glutamatergic and non-glutamatergic axon terminals.

**Three Key Points:**

1. A direct anterograde projection from orexin-A-expressing neurons in the hypothalamus to the mouse olfactory bulb (OB) was confirmed using virally mediated tract tracing.
2. Orexin-A and melanin-concentrating hormone (MCH), two neuropeptides primarily produced in the lateral hypothalamus are broadly expressed throughout the mouse OB.
3. Approximately 64% of putative orexin-A axon terminals overlapped with vesicular glutamate transporter type 1 (VGLUT1) or type 2 (VGLUT2). In contrast, only 25% of putative MCH axon terminals overlapped with VGLUT2.

## INTRODUCTION

In principle, neural circuits that can adjust olfactory sensory processing based on an organism’s internal state would facilitate perceptual functions such as recognizing and locating food, mates, and threats (Gross-Isseroff & Lancet, 1988; Homma, Cohen, Kosmidis, & Youngentob, 2009; Rokni, Hemmelder, Kapoor, & Murthy, 2014; Uchida & Mainen, 2007). A current model proposes that state-dependent modulation can occur in the olfactory bulb (OB), the primary site that receives input from olfactory receptor neurons (Chelette et al., 2021; Fardone et al., 2019; Julliard, Al Koborssy, Fadool, & Palouzier-Paulignan, 2017; Kolling et al., 2022). The OB can transform sensory input based on a local synaptic network and centrifugal inputs from other brain regions, which includes a direct projection from the lateral hypothalamus (In’t Zandt, Cansler, Denson, & Wesson, 2019; Qi, Fadool, & Storace, 2023; Schneider et al., 2020; Shipley & Adamek, 1984). The lateral hypothalamus is a brain area involved in homeostatic regulation and is a candidate mechanism to link sensory processing with state-dependent signaling (Berthoud & Münzberg, 2011; Fu et al., 2019; Timper & Bruning, 2017; Yamanaka et al., 2003).

The direct projections from the lateral hypothalamus to the OB originate from a genetically heterogeneous population of neurons, of which approximately 22% express the neuropeptide orexin-A (Qi et al., 2023). The neurons that did not express orexin-A (∼78%) originated from a region of the lateral hypothalamus that is known to contain neurons that express the neuropeptide melanin-concentrating hormone (MCH) (Broberger, De Lecea, Sutcliffe, & Hökfelt, 1998; Qi et al., 2023; Y. Saito & Nagasaki, 2008; Schneider et al., 2020; Wheeler et al., 2014). Orexin-A and MCH neurons are both involved in functions that include sleep, motivation, metabolism and ingestive behaviors (Aston-Jones et al., 2010; Concetti, Peleg-Raibstein, & Burdakov, 2024; Dawson et al., 2023; Diniz & Bittencourt, 2017; Peyron et al., 2000; Subramanian et al., 2023; Thannickal et al., 2000). However, no study to date has reported the presence of MCH expression in the mouse OB (Bittencourt et al., 1992; Croizier et al., 2013; Y. Saito, Cheng, Leslie, & Civelli, 2001; Schneider et al., 2020).

Additionally, it is unclear whether the OB is broadly or selectively innervated by the projections from the lateral hypothalamus. We addressed these questions using virally mediated anterograde tract tracing and immunohistochemistry for orexin-A and MCH. Orexin-A and MCH expression, including their putative axon terminals, were present throughout the OB with similar densities in all anatomical layers from anterior to posterior. These data confirmed previous reports of orexin-A expression in the rat OB and provide the first report of MCH expression in the mouse OB (Caillol, Aïoun, Baly, Persuy, & Salesse, 2003; Hardy et al., 2005; Shibata et al., 2008).

Both orexin-A and MCH-expressing neurons co-express at least one VGLUT isoform, which are proteins that define different classes of glutamatergic neurons (Chee, Arrigoni, & Maratos-Flier, 2015; El Mestikawy, Wallén-Mackenzie, Fortin, Descarries, & Trudeau, 2011; Fremeau et al., 2001; Henny, Brischoux, Mainville, Stroh, & Jones, 2010; Herzog et al., 2001; Mickelsen et al., 2017; Pham et al., 2024; Rosin, Weston, Sevigny, Stornetta, & Guyenet, 2003; Schneeberger et al., 2018; Schöne, Apergis-Schoute, Sakurai, Adamantidis, & Burdakov, 2014; Schöne et al., 2012; Takamori, Rhee, Rosenmund, & Jahn, 2000). It’s unknown whether the orexin-A and MCH-expressing axon terminals that innervate the mouse OB are themselves genetically heterogeneous.

We addressed this question using high-resolution confocal microscopy to quantify whether orexin-A and MCH-expressing axon terminals also co-express VGLUT1 or VGLUT2. We used the morphology of orexin-A and MCH expression to identify putative axon terminals originating from both populations of neurons. Nearly half of the orexin-A-expressing axon terminals overlapped with VGLUT2 alone (39%), while smaller proportions overlapped with VGLUT1 alone (6%), both VGLUT1 and VGLUT2 (18%) or did not overlap with either VGLUT (23%). In comparison, 25% of MCH-expressing axon terminals overlapped with VGLUT2, with the majority (72%) not overlapping with either VGLUT1 or VGLUT2.

Therefore, the hypothalamic projections to the mouse OB include at least two neurochemically distinct subpopulations, each of which are genetically heterogeneous. Additionally, our results indicate that the projections from both orexin-A and MCH-expressing neurons to the OB are capable of releasing glutamate. This study supports a model in which the lateral hypothalamic projections to the mouse OB can modulate olfactory sensory processing in a neurochemically heterogeneous manner.

## MATERIALS AND METHODS

### Solutions, reagents, and equipment

**TABLE 1.** List of all solutions, reagents, and equipment used in this study.

| Solution/reagent | Source | Product / RRID # |
| --- | --- | --- |
| Mice (C57BL/6J) | Jackson Laboratories | 000664 |
| Ketamine hydrochloride | Zoetis | 010177 |
| Anased | Akorn | 033197 |
| Sodium chloride | Hospira | 061758 |
| Atropine sulfate | Covetrus | 074760 |
| Dexamethasone | Bimeda | 069238 |
| Carprofen | Zoetis | 024751 |
| Bupivacaine hydrochloride | Hospira | 071286 |
| Nair | Church and Dwight |  |
| Paralube Vet Ointment | Dechra Veterinary Products |  |
| Euthasol | Virbac | 009444 |
| Stereotax | David Kopf | Model 963 |
| DC Temperature Controller | FHC | 40-90-8D |
| Dental drill | Osada | XL-230 |
| Sutures | Covetrus | 041178 |
| Nanoject III | Drummond Scientific | 3-000-207 |
| Glass micropipettes | Drummond Scientific | 3-000-203-G/X |
| P-97 Pipette puller | Sutter Instruments | RRID:<br>SCR_016842 |
| VT1000S Vibratome | Leica | RRID:<br>SCR_016495 |
| Phosphate-buffered saline | MP Biomedicals | 2810301 |
| rAAV-Hypocretin-EGFP-WPRE-hGH polyA | BrainVTA | PT5293 |
| Paraformaldehyde | Electron Microscopy Services | 15714-S |
| TWEEN20 | Sigma–Aldrich | 9005-96-5 |
| Triton-X-100 | Sigma–Aldrich | 9036-19-5 |
| Sodium borohydride (NaBH<sub>4</sub>) | BeanTown Chemical | 125800-100G |
| TrueBlack | Biotium | 23007 |
| Normal Goat Serum | ThermoFisher Scientific | 31872 |
| Rabbit anti-orexin-A primary antibody (Ab6214) | Abcam | RRID: AB_305380 |
| Mouse anti-orexin-A primary antibody (Ab89886) | Abcam | Ab89886 |
| Rabbit anti-MCH primary antibody (Ab274415) | Abcam | RRID: AB_3101761 |
| Mouse anti-VGLUT2 primary antibody (135421) | Synaptic Systems | RRID: AB_2884916 |
| Guinea pig anti-VGLUT1 primary antibody (135304) | Synaptic Systems | 135304 |
| iFluor 488 goat anti-rabbit | AAT Bioquest | ABD-16678 |
| Alexa Fluor Plus 555 goat anti-rabbit | Invitrogen | A32732 |
| iFluor 633 goat anti-rabbit | AAT Bioquest | 16638 |
| iFluor 488 goat anti-mouse | AAT Bioquest | 16528 |
| Alexa Fluor Plus 555 goat anti-mouse | Invitrogen | A32727 |
| Alexa Fluor 647 goat anti-guinea pig | Abcam | ab150187 |
| Recombinant Human PMCH protein | Abcam | Ab314595 |
| Fluoromount-G mounting medium with DAPI | SouthernBiotech | 0100-20 |
| Stereo Investigator | MBF Biosciences | RRID:<br>SCR_002526 |
| Nikon Confocal (microscope) | Nikon Instruments | CSU-W1 |
| Nikon NIS-Elements (software) | Nikon Instruments | RRID:<br>SCR_014329 |
| MATLAB (software) | Mathworks | RRID:<br>SCR_001622 |
| Adobe Creative Cloud (software) | Adobe Systems | RRID:<br>SCR_010279 |
| ImageJ (software) | National Institutes of Health | RRID:SCR_003070 |

### Animals and Ethics Statement

All experimental procedures were performed in accordance with institutional requirements approved by the Florida State University (FSU) Animal Care and Use Committee. Our study used adult (6-24 weeks of age) male (N = 15) and female (N = 8) C57BL/6J mice that were housed in the FSU animal vivarium with a 12/12-hour light/dark cycle and ad libitum access to food and water.

### Stereotaxic Viral Injection

Mice were anesthetized with ketamine/xylazine (90/10 mg/kg) and were given pre-operative injections of atropine (0.2 mg/kg), dexamethasone (4 mg/kg), and carprofen (20 mg/kg). Anesthetized mice were maintained on top of a heating pad (DC Temperature Controller; FHC) until regaining sternal recumbency. Anesthetic depth was regularly monitored via pedal reflex and additional anesthesia was administered as needed to ensure that the mice remained anesthetized during the procedure. After reaching a stable plane of anesthesia, mice were administered eye ointment, and the hair on top of the head was removed using a depilatory (Nair; Church and Dwight) followed by multiple saline rinses. Mice were placed in a stereotaxic device (Model 963; David Kopf Instruments) and the skin over the skull was cleaned and disinfected using iodine and alcohol (Covidien).

Bupivacaine (2 mg/kg) was injected subcutaneously before making an incision.

A craniotomy was made over the right hypothalamus (1.5-1.7 mm posterior to bregma and 1 mm lateral to the sagittal fissure) using a dental drill (Osada XL-230). A glass micropipette with a tip diameter of 10-20 μm was made using a P-97 micropipette puller and was filled with AAV2/9-Hypocretin-EGFP-WPRE-hGH-polyA (Brain VTA, PRODUCT # PT5293). The three mice received up to 1000 nl of total injection volumes across two anterior-posterior sites (1.5 mm and 1.7 mm posterior to Bregma) within the craniotomy separated by 0.2 mm. For each injection site the filled micropipette was lowered into the right hypothalamus using a stereotaxic arm (1 mm lateral to the sagittal fissure and 5.2 mm ventral). The virus was delivered in multiple volumes of 100 nL at a rate of 4 nL/s, with a minimum of a 2-minute wait between additional injections. The glass pipette was left in place for at least 2 minutes to facilitate viral diffusion after each injection. After the micropipette was removed from the brain, the incision was sutured (polyglycolic acid braided absorbable sutures, Covetrus), and the animals were monitored until regaining sternal recumbency, after which they were returned to the vivarium. Animals received a second postoperative dose of carprofen at the end of the day of surgery, and for the first 3 days after the procedure.

### Immunohistochemistry

After allowing the virus to express for 14 days, mice were anesthetized by injections of ketamine/xylazine and Euthasol and then were perfused transcardially with PBS and 4% paraformaldehyde in PBS. After perfusion, the brain was removed, postfixed overnight in 4% paraformaldehyde solution at 4°C, and cut into 40-μm-thick coronal sections on a vibratome (Leica VT1000 S).

The immunolabeling procedure was conducted as follows: free floating sections were washed in 1% NaBH_4_ in PBS for 20 minutes to reduce background autofluorescence, followed by 3 washes in PBST (0.1% Tween20 in PBS) for 10 minutes each, and then were washed with PBSTT (0.1% Triton X-100 in PBST) for 20 minutes. Lipofuscin autofluorescence was quenched by incubating the sections in 1X TrueBlack (Biotium, diluted from 20X TrueBlack stock solution in 70% ethanol) for 30 seconds, followed by 3 washes with PBST for 5 minutes each. Sections were then incubated in blocking solution (5% normal goat serum in PBST) for 30 minutes and then were incubated in primary antibody solution (1:1000, diluted in blocking solution) overnight at 4°C. The next day, sections were washed 3 times in PBS for 5 minutes each, incubated with the secondary antibody (1:500, diluted in PBST) for 1 hour at room temperature, and then were washed 3 times in PBS for 5 minutes each.

The sections were mounted on clean slides with DAPI Fluoromount-G mounting medium applied to the coverslips (5-10 drops per slide). Solutions of 1% NaBH_4_ in PBS, 1X TrueBlack, blocking solution, primary and secondary antibody were made fresh on the day of the experiment. The combinations of primary and secondary antibodies for different immunostaining reactions are listed in Table 2.

**TABLE 2.** Combinations for immunostaining.

| Series | Primary antibody | Secondary antibody |
| --- | --- | --- |
| Overlap of orexin-A and eGFP expression induced by AAV-orexin | Rabbit anti-orexin-A | Goat anti-rabbit Alexa Fluor Plus 555 |
| Orexin-A expression | Rabbit anti-orexin-A | Goat anti-rabbit iFluor 488 |
| Validation of MCH antibody | Rabbit anti-MCH | Goat anti-rabbit Alexa Fluor Plus 555 |
| MCH expression | Rabbit anti-MCH | Goat anti-rabbit iFluor 488 |
| Validation of mouse anti-orexin-A antibody | Mouse anti-orexin-A<br>Rabbit anti-orexin-A | Goat anti-mouse iFluor 488<br>Goat anti-rabbit iFluor 633 |
| Co-labeling of orexin-A and MCH | Mouse anti-orexin-A<br>Rabbit anti-MCH | Goat anti-mouse iFluor 488<br>Goat anti-rabbit iFluor 633 |
| Co-labeling of orexin-A and VGLUT2 | Rabbit anti-orexin-A<br>Mouse anti-VGLUT2 | Goat anti-rabbit iFluor 488<br>Goat anti-mouse Alexa Fluor Plus 555 |
| Triple staining of orexin-A, VGLUT1, and VGLUT2 | Rabbit anti-orexin-A<br>Guinea pig anti-VGLUT1<br>Mouse anti-VGLUT2 | Goat anti-rabbit iFluor 488<br>Goat anti-guinea pig Alexa Fluor 647<br>Goat anti-mouse Alexa Fluor Plus 555 |
| Triple staining of MCH, VGLUT1, and VGLUT2 | Rabbit anti-MCH<br>Guinea pig anti-VGLUT1<br>Mouse anti-VGLUT2 | Goat anti-rabbit iFluor 488<br>Goat anti-guinea pig Alexa Fluor 647<br>Goat anti-mouse Alexa Fluor Plus 555 |

The specificity of the rabbit anti-MCH primary antibody (Catalog # ab274415; Abcam) was confirmed in a preincubation study. The MCH primary antibody solution (1:1000 diluted in blocking solution) was preincubated overnight at 4°C with PMCH (Abcam, Lot # ab314595) at a concentration of 5.5 μg/μL.

### Confocal imaging

All images in this study were acquired using a Nikon CSU-W1 spinning disk confocal microscope equipped with 405 nm, 488 nm, 561 nm, and 640 nm laser lines. DAPI and eGFP were excited using the 405 nm and 488 nm laser lines, respectively. Secondary antibodies conjugated to 488 nm, 555 nm and 633 nm dyes were excited using the 488 nm, 561, and 640 nm laser lines, respectively.

### Generation of immunohistochemical plots

Large tiled confocal images were acquired using a 20x 0.95 N.A. objective lens in 2 μm z-steps through the depth of sections with the large image function in NIS elements software (Nikon Instruments Inc) (**Figure 3a, 3d**). The images of individual processes in were cropped from these large, tiled images and were individually adjusted for brightness and contrast in Fiji (**Figure 3a, d**, 1-4). Orexin-A and MCH expression was quantified in maximum z-projections of the OB images using the Labkit plugin in Fiji (**Figure 3b, e**) (Arzt et al., 2022). The Labkit classifier was trained by the experimenter by marking between 5-10 immunoreactive markers and 5-10 locations containing background signal. The same classifier was applied to the entire dataset for pixel-level segmentation. The quality of the segmentation was visually screened in each section by the experimenters, and the classifier underwent additional training when necessary. The resulting segmentation was displayed in Fiji, and segmented areas greater than 2 pixels (0.845 μm^2^) were identified using the ‘Analyze Particles’ function. The contour lines of each OB layer were manually traced in Fiji using DAPI fluorescence, and the bregma position and neuroanatomical structures were identified by aligning the DAPI fluorescence images with a mouse brain atlas (Franklin & Paxinos, 2008). The resulting segmentation data and drawn contours were imported into MATLAB (Mathworks) via a custom script in which label density was quantified for each layer. The density values were calculated as the number of segmented regions of interest divided by the area of each OB layer.

### Morphology and co-expression analysis

Size and co-expression analyses of orexin-A and MCH labeled processes were performed in high-resolution confocal images in which sections were imaged using a 100x 1.45 N.A. objective lens in 0.2 μm z-steps through each section. Qualitative assessment of orexin-A and eGFP overlap was conducted in 9 images across 3 different preparations. For quantification of orexin-A and MCH labeled processes, the confocal images were imported into Stereo Investigator (Stereo Investigator, Microbrightfield) software and orexin-A or MCH expression was manually labeled. For every clearly labeled neuropeptide process, the largest cross-sectional area within a z-projection up to 2 μm was identified and measured using the nucleator probe in Stereo Investigator. The center of each area was intersected with 6 equally spaced lines, and the distance from the center to the edge of each process was marked. The area of each process was calculated as A = π × r^2^, where r is the mean of the distances from the center to the edge of each marker. To quantify overlap with VGLUTs, co-expression of orexin-A or MCH immunoreactive product with VGLUT1 and VGLUT2 was identified based on the presence of spatially overlapping fluorescence signals that exhibited similar morphology in the z-projection.

### Statistical analysis

All statistical analyses were performed on MATLAB. Comparisons between groups were conducted using an N-way ANOVA. Post-hoc pairwise comparisons were performed using the Tukey’s honestly significant difference (HSD) test to identify specific group differences. A p-value < 0.05 was considered statistically significant, with results annotated as follows: ns, no significant difference; *, p < 0.05; **, p < 0.01; ***, p < 0.001. Data are presented as mean ± standard error of the mean (SEM).

## RESULTS

### Anterograde projections from orexin-A neurons to the mouse OB

We confirmed the presence of a direct anterograde projection from orexin-A-expressing neurons in the lateral hypothalamus to the mouse OB by injecting an adeno-associated virus (AAV) that expressed eGFP under the control of the hypocretin (hereafter referred to as orexin) promoter into the right hypothalamus (**Figure 1a-b**, rAAV2/9-Hypocretin-EGFP-WPRE-hGH polyA, N = 3 different mouse preparations). eGFP expression was present in neurons and their peripheral dendrites located in the lateral hypothalamus, the brain area known to contain orexin-A-expressing neurons (Gascuel et al., 2012; Nambu et al., 1999; Peyron et al., 1998; Sakurai et al., 1998) (**Figure 1c**, eGFP). eGFP label was absent in the somata of neurons located in the contralateral hypothalamus (not shown).

**Figure 1:**
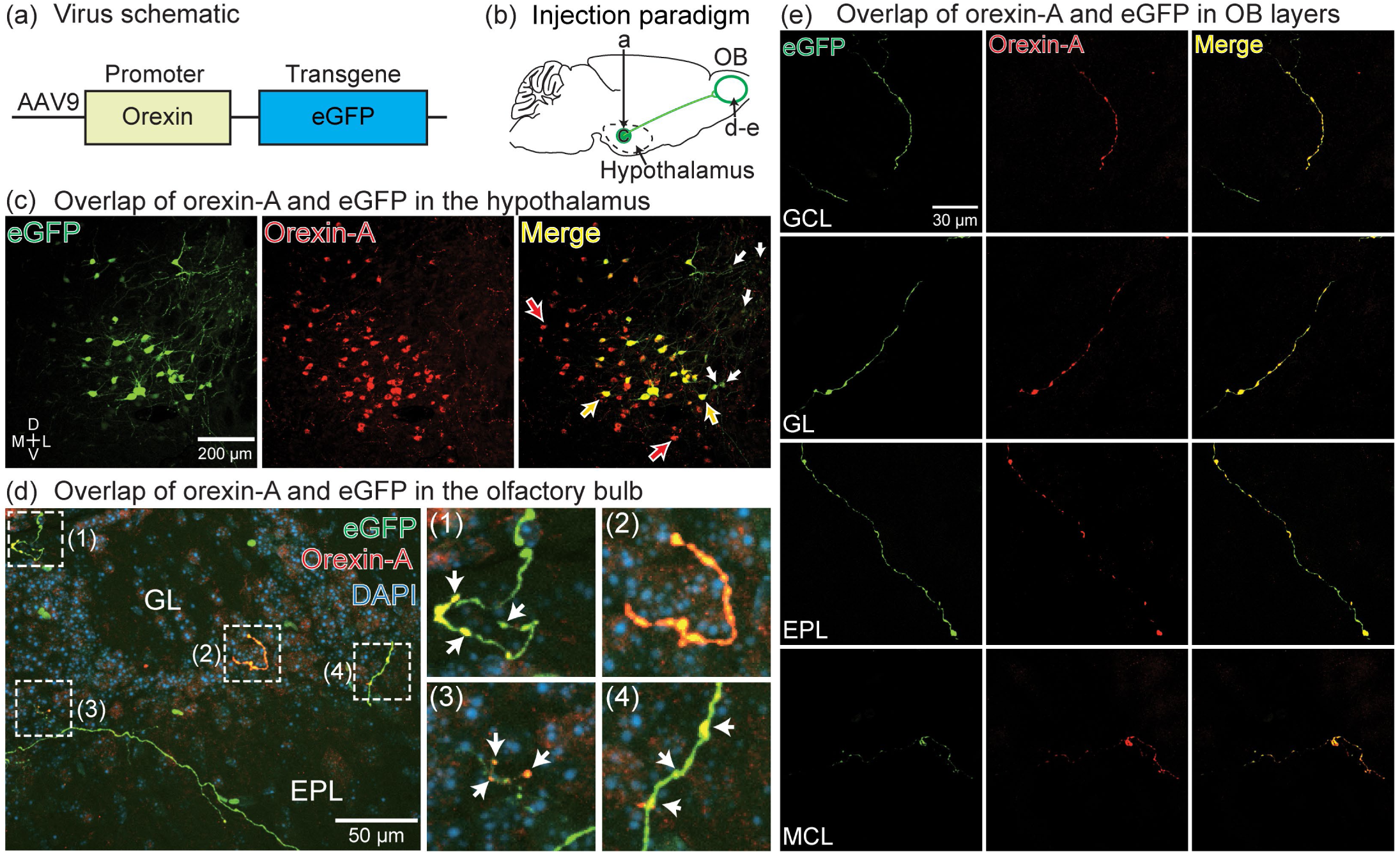
(a-b) Schematic of the orexin-eGFP AAV and the injection paradigm into the hypothalamus. (c) Histology through the hypothalamus illustrating eGFP and orexin-A expression. The orange arrows point to neurons with overlapping eGFP and orexin-A. The white arrows point to eGFP-expressing neurons that do not overlap with orexin-A (5/50 neurons in this section). The red arrows point to orexin-A neurons that did not express eGFP. (d) eGFP (green) and orexin-A (red) expression in the OB illustrating co-expression in the anterograde projections across multiple OB layers. Magnified insets (1-4) reveal fine details of eGFP and orexin-A overlap in axon varicosities (white arrows). (e) Additional examples illustrating eGFP-expressing processes and their overlap with orexin-A in different OB layers. GCL, granule cell layer; GL, glomerulus layer, EPL, external plexiform layer, MCL, mitral cell layer.

We performed immunohistochemistry for orexin-A to evaluate the ability of the orexin promoter to drive eGFP expression in orexin-A-expressing neurons. Orexin-A-immunoreactive neurons were prominently located in the lateral hypothalamus, consistent with prior reports (**Figure 1c**, Orexin-A) (Nambu et al., 1999; Peyron et al., 1998; Qi et al., 2023; Sakurai et al., 1998). Most of the eGFP-expressing neurons overlapped with orexin-A-immunoreactive product (hereafter referred to as orexin-A expression) (**Figure 1c**, Merge). The proportion of eGFP-expressing neurons that overlapped with orexin-A expression was quantified in the 3 sequential sections spanning 480 µm through the lateral hypothalamus that contained the most eGFP-expressing neurons in each preparation (583 eGFP-expressing cells in 3 mouse preparations; 176-223 neurons per preparation). Orexin-A overlapped with eGFP in 86.6 ± 16.2% of the eGFP-expressing neurons (**Figure 1c**, Merge, orange arrows; the proportions ranged between 67.4% - 96.9%). eGFP-expressing neurons that overlapped with orexin-A exhibited qualitatively stronger eGFP fluorescence than the small number of eGFP-expressing neurons that did not overlap with orexin-A (**Figure 1c**, Merge, the white arrows point to the eGFP-expressing neurons that did not overlap with orexin-A). Many orexin-A-expressing neurons did not overlap with eGFP, which is consistent with the eGFP expression being dependent on the location of the AAV injection (**Figure 1c**, Merge, red arrows). eGFP expression was present in axon-shaped processes in different layers throughout the OB, and often (but not always) overlapped with orexin-A expression (**Figure 1d-e**). Processes containing eGFP expression were present in other brain areas, but not analyzed here (e.g., cortex, hippocampus). Therefore, AAV transduction using the orexin promoter can drive eGFP expression into orexin-A neurons, and orexin-A expression overlaps with the axon projections of orexin-A neurons in the mouse OB.

### Melanin-concentrating hormone (MCH) is expressed in the mouse OB

Most of the lateral hypothalamic projections to the OB originate from neurons that do not express orexin-A (Qi et al., 2023). We hypothesized that this pathway includes neurons that express the neuropeptide melanin-concentrating hormone (MCH), which are located near neurons that express orexin-A (Broberger et al., 1998; Mickelsen et al., 2017; Qi et al., 2023; Qu et al., 1996; Rao et al., 2008; Schneider et al., 2020). Immunohistochemistry using a rabbit anti-MCH antibody labeled neurons in the lateral hypothalamus (**Figure 2a-b**). We validated the specificity of the MCH antibody by performing immunohistochemistry on adjacent sections after it had been preincubated overnight with MCH. Sections that underwent treatment with the preincubated antibody contained no labeled neurons or processes (**Figure 2c**, similar results were observed in sections from 2 mouse preparations).

**Figure 2:**
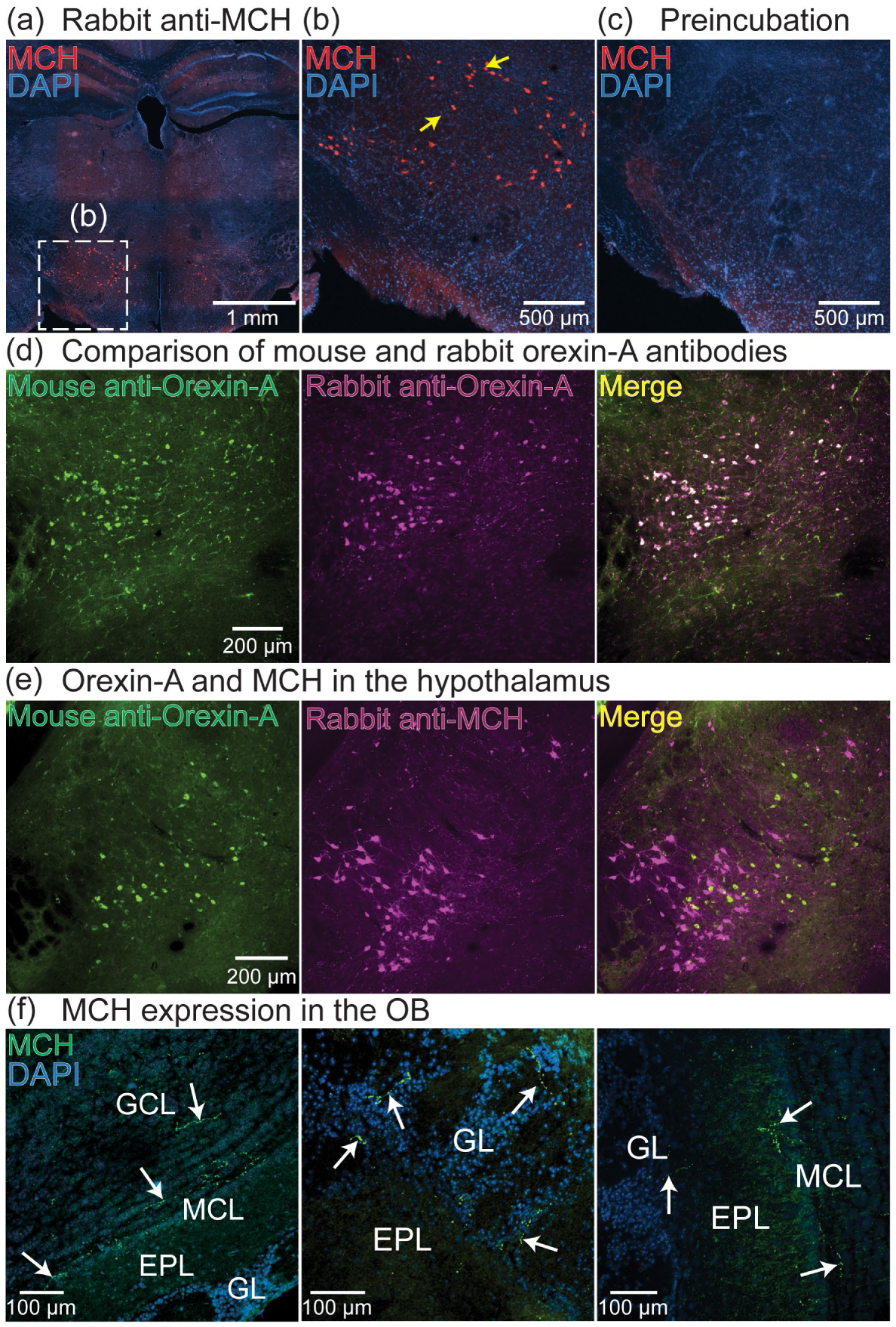
(a-b) Low (a) and high-magnification (b) images of MCH-expressing neurons in the lateral hypothalamus. Yellow arrows indicate labeled neurons. (c) An adjacent section from the same preparation in panels a-b that was processed using the rabbit anti-MCH primary antibody after it had been preincubated overnight with MCH. (d) Overlapping expression patterns from two different orexin-A primary antibodies in the lateral hypothalamus. (e) Orexin-A and MCH-expressing neurons in the lateral hypothalamus labeled in the same section. (f) MCH expression in the mouse OB.

**Figure 3:**
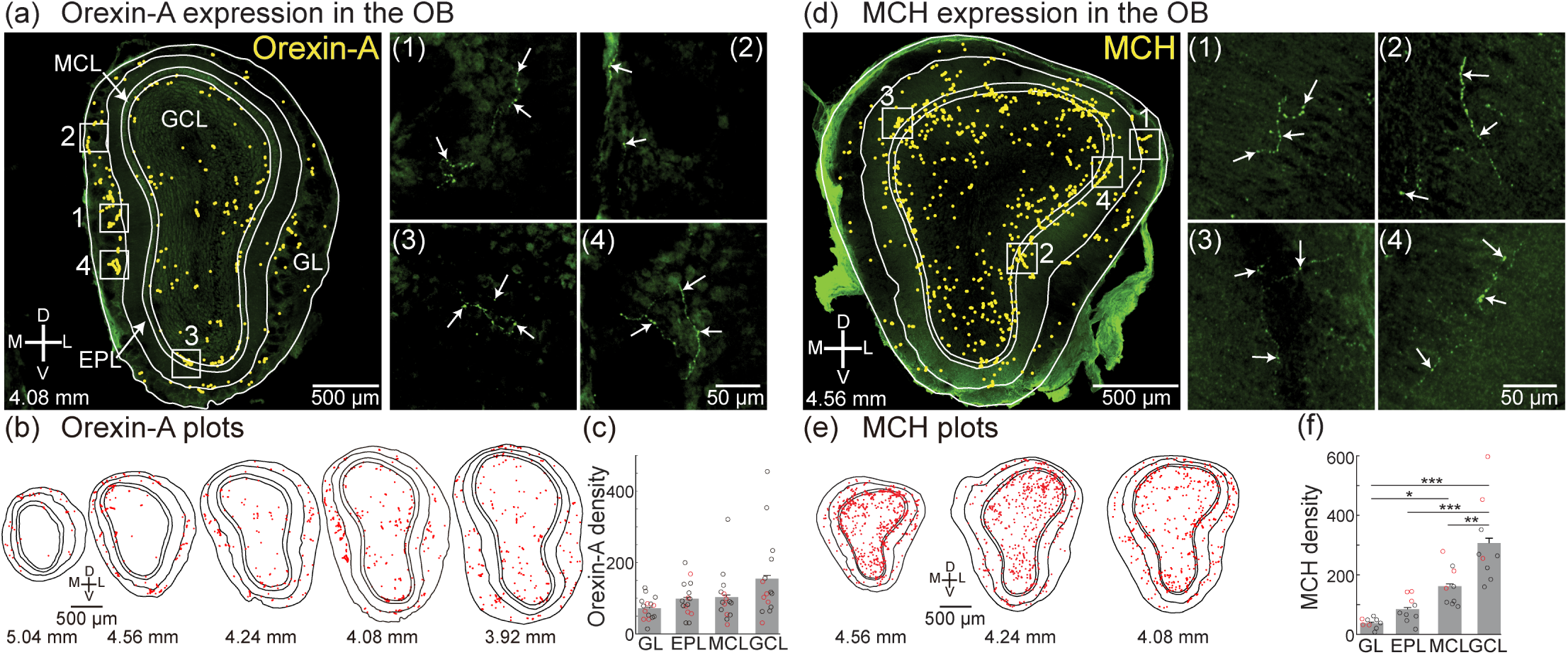
(a, left) Tiled confocal image of an entire OB section with layers outlined in white and yellow dots that indicate the presence of orexin-A expression. (a, 1-4) Cropped images from subpanel a illustrating orexin-A expression in the corresponding white boxes. (b) Plots of orexin-A expression across sequential OB sections at different anterior-posterior positions from one preparation. Each marker represents an orexin-A immunoreactive product with a minimum size of 0.845 μm^2^. (c) Orexin-A density in each OB layer. Each circle represents the mean density from an individual preparation (the red circles are from the preparation in subpanels a-b). (d-f) Same layout as panels a-d but for MCH.

Because orexin-A and MCH are expressed in non-overlapping cell populations within the lateral hypothalamus, we further validated the specificity of the MCH primary antibody by comparing their label patterns in the same sections. We first identified and validated a primary antibody for orexin-A that was raised in a host species (mouse-anti-orexin-A) that was compatible with the rabbit anti-MCH antibody used in **Figure 2a-c**. We validated the mouse anti-orexin-A antibody by comparing its labeling pattern to that of a previously validated rabbit anti-orexin-A antibody (**Figure 2d**, left panel) (Qi et al., 2023). Sections that underwent immunohistochemistry with both mouse anti-orexin-A and rabbit anti-orexin-A antibodies had completely overlapping labeled neurons in the lateral hypothalamus (**Figure 2d**, similar results were obtained in 3 different mouse preparations, all labeled neurons overlapped). Sections that underwent immunohistochemistry using both the mouse-anti-orexin-A and rabbit anti-MCH antibodies contained two non-overlapping populations of labeled neurons in the lateral hypothalamus (**Figure 2e**). Therefore, the rabbit anti-MCH primary antibody specifically labels the MCH neuron population. Immunohistochemistry performed on OB sections using the rabbit anti-MCH antibody revealed the presence of MCH expression in different layers throughout the OB (**Figure 2f**, white arrows).

### Orexin-A and MCH are broadly expressed throughout the mouse OB

We quantified the spatial distribution of orexin-A and MCH expression in sequential sections from anterior-to-posterior (Bregma 5.04 mm to 3.92 mm) through the mouse OB in large tiled confocal images acquired in 2 µm z-steps through the entire section using a 20x 0.95 N.A. lens (**Figure 3a**, quantification was done in 3 different mouse preparations). This approach provided the spatial resolution needed to visualize individual processes throughout entire sections (**Figure 3a**, right subpanels). Orexin-A was expressed in the OB at a mean density of 119.6 ± 63.9 / mm^2^ (N = 15 sections, ranged from 49.3 products/mm^2^ to 297.2 products/mm^2^). Orexin-A expression was present in each layer within the OB, with no significant differences in its density across the layers (GCL: 154.9 ± 30 products/mm^2^; MCL: 103.4 ± 18.2 products/mm^2^; EPL: 98.5 ± 12.4 products/mm^2^; GL: 72 ± 8.2 products/mm^2^) (**Figure 3b-c**).

We used the same approach to quantify the MCH expression pattern within the OB, which was measured at a mean density of 171.2 ± 83.5/mm^2^ (**Figure 3d-f**, N = 3 mouse preparations, 3 sections per preparation; 90.1 – 307.9 labeled processes/mm^2^). MCH expression was present in all OB layers and along the anterior-posterior axis, with significantly highest densities present in the granule cell layer (GCL: 306.4 ± 46.7 labeled processes/mm^2^; MCL: 161.4 ± 21.9 products/mm^2^; EPL: 84.4 ± 14.4 labeled processes/mm^2^; GL: 38 ± 5.6 labeled processes/mm^2^) (**Figure 3e-f**). Therefore, the OB is broadly innervated by both orexin-A-and MCH-expressing processes, although MCH expression is more spatially organized with respect to anatomical layers.

### Identifying putative axon terminals from orexin-A and MCH-expressing neurons in the OB

The expression of both orexin-A and MCH was observed as fine processes and larger bulbous varicosities. To test whether the morphology of orexin-A expression in the OB reflects axon terminals or fibers of passage, we quantified the size of every clearly labeled orexin-A process in high resolution (100x 1.45 N.A.) confocal images. The mean area of all orexin-A processes was 1.31 ± 0.98 µm^2^, which ranged from 0.18 to 4.76 µm^2^ (N = 498 orexin-A-expressing processes measured in 6 sections from 3 different preparations) (**Figure 4a**). Sorting the size distribution of orexin-A labeled processes from smallest to largest within individual confocal images revealed graded increases and occasional sharp transitions (**Figure 4b**, points reflect the distribution of sizes from one exemplar field of view). We quantified this transition point by plotting the derivative of adjacent markers in the sorted size distributions (**Figure 4c**). We defined the maximum value in the derivative plot as the peak (at least 3 times the mean of the derivative), and defined the midpoint of the two points that generate this peak as the size transition point between fibers of passage and axon terminals, which ranged between 0.72 - 0.99 µm^2^ (**Figure 4d**, each point is the size transition point from an individual confocal image). We performed the same analysis in 26 images of MCH-expressing processes in OB sections from 3 different mouse preparations (8-9 images from different sections per preparation). MCH expression ranged in size from 0.07 to 3.1 µm^2^ (mean of 0.65 ± 0.37 µm^2^; N = 2,259 MCH-expressing products) (**Figure 4e**). Using the same morphological analysis as for orexin-A-expressing products, we determined a size threshold for MCH putative axon terminals to be 0.73 ± 0.11 µm^2^ (thresholds ranged between 0.49 - 0.86 µm^2^) (**Figure 4f-h**). Orexin-A and MCH expression with an area greater than 0.88 µm^2^ and 0.73 µm^2^, respectively, are hereafter referred to as putative axon terminals. Notably, the orexin-A putative axon terminals were significantly larger than those from the MCH-expressing population (p < 0.001).

**Figure 4:**
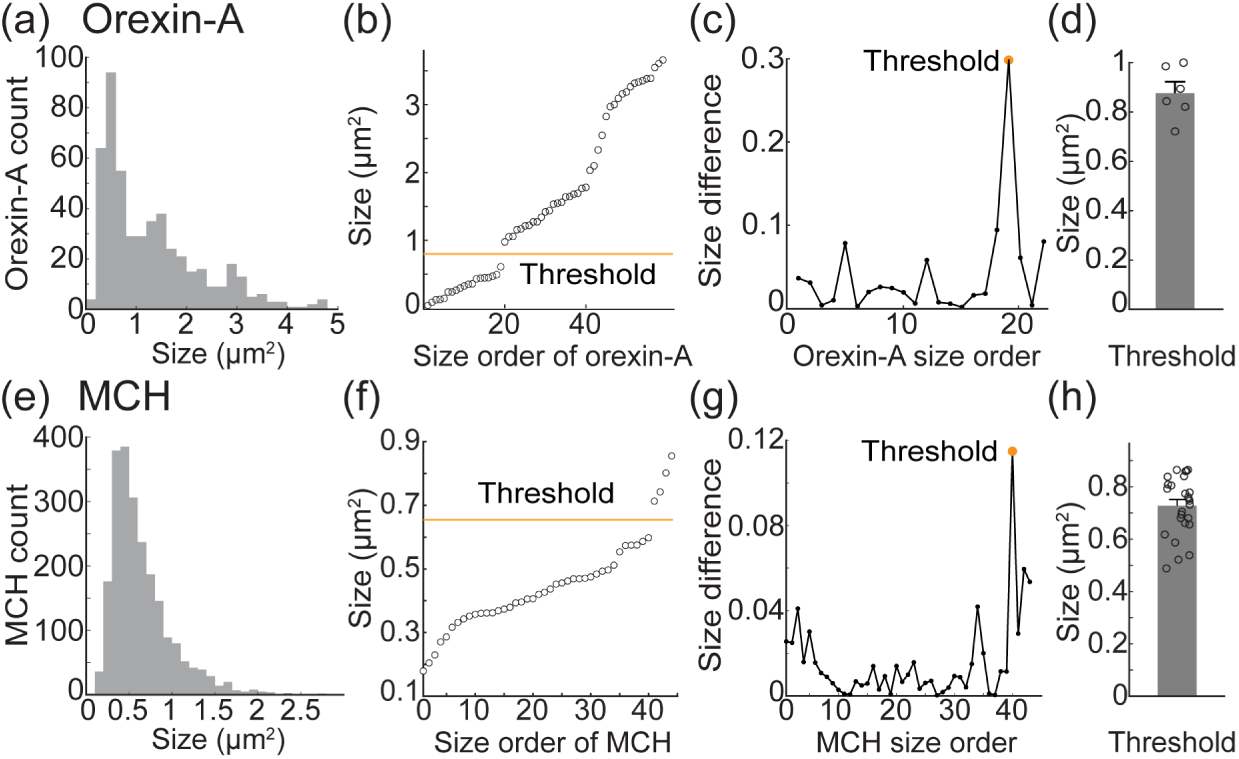
(a) Size of all individual orexin-A labeled processes in the dataset. (b) The size of all orexin-A processes in a single section sorted from smallest to largest. (c) The difference in label size in the neighboring markers from panel c. The yellow marker indicates the size threshold used to differentiate between small and large varicosities and is plotted on panel b. (d) Mean size used to define putative axon terminals. (e-h) Same layout as panels a-d but for MCH.

### Glutamatergic heterogeneity within orexin-A and MCH axon terminals in the OB

Orexin-A-and MCH-expressing neurons are genetically heterogeneous, with different subsets that express vesicular glutamate transporter type 1 (VGLUT1) or type 2 (VGLUT2) (Chee et al., 2015; El Mestikawy et al., 2011; Fremeau et al., 2001; Henny et al., 2010; Herzog et al., 2001; Mickelsen et al., 2017; Pham et al., 2024; Rosin et al., 2003; Schneeberger et al., 2018; Schöne et al., 2014; Schöne et al., 2012; Takamori et al., 2000). The OB contains complementary distributions of VGLUT1 and VGLUT2 (**Figure 5**) (Gabellec, Panzanelli, Sassoè-Pognetto, & Lledo, 2007; Nakamura, Hioki, Fujiyama, & Kaneko, 2005; Ohmomo et al., 2009). VGLUT2 expression is most prominent in the glomerular layer, with lighter expression in the mitral and granule cell layers (**Figure 5**, VGLUT2). In contrast, VGLUT1 is weakly expressed in the glomerular layer, with dense expression in the external plexiform and granule cell layers (**Figure 5**, VGLUT1). To test whether the OB projections from orexin-A and MCH-expressing neurons reflect their glutamatergic heterogeneity, we quantified the overlap of their putative axon terminals with VGLUT1 and VGLUT2.

**Figure 5:**
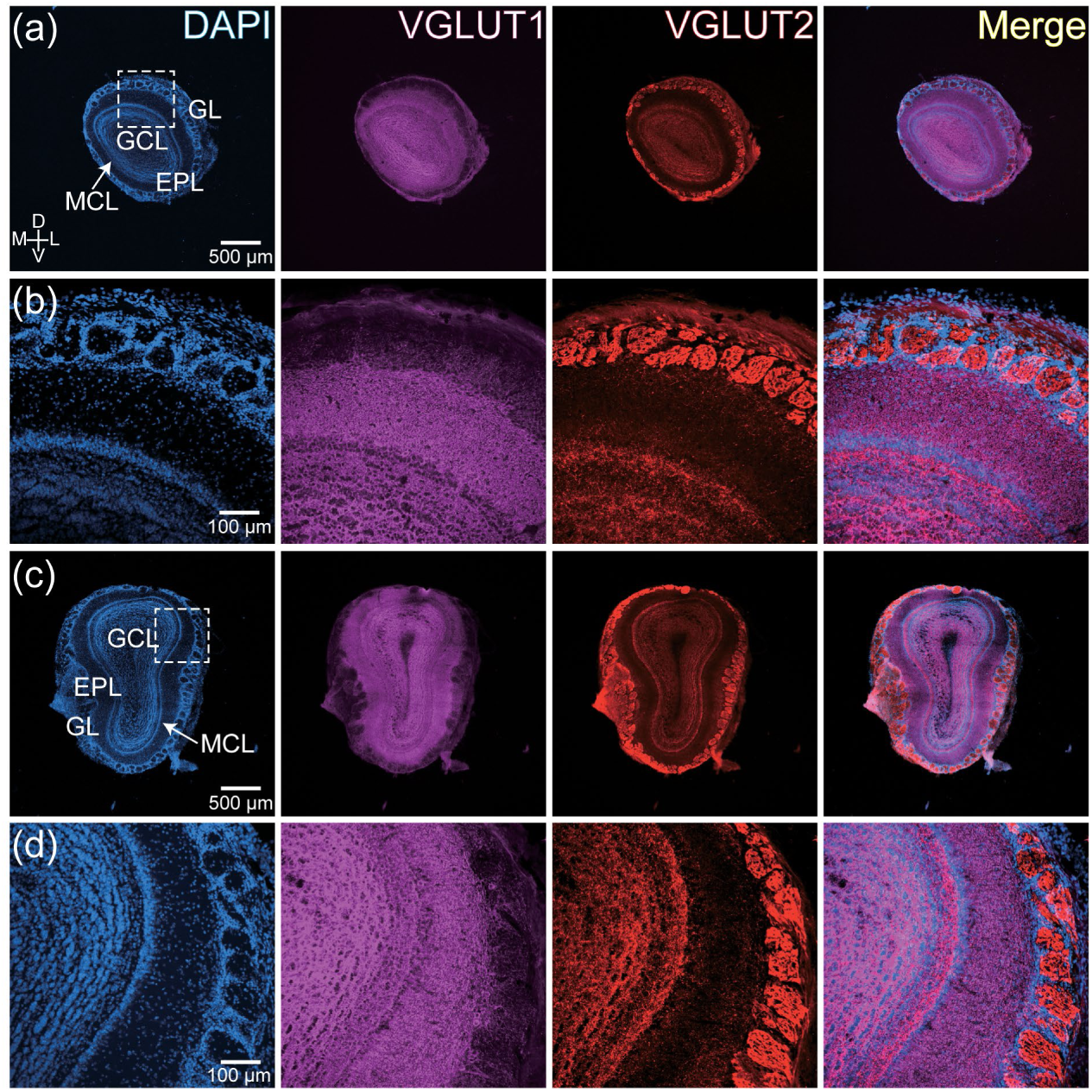
(a-b) Low (a) and high (b) magnification images of the anterior OB illustrating DAPI (blue), VGLUT1 (magenta) and VGLUT2 (red) label. Subpanel b is taken from the boxed area in subpanel a. (c-d) Same arrangement as subpanels a-b for a section from the posterior OB.

Overlap between orexin-A putative axon terminals, VGLUT1 and VGLUT2 was quantified in 30 confocal images taken from different OB layers in 3 preparations (8-12 images per preparation, N = 1205 terminals) (**Figure 6**). Orexin-A axon terminals that did not overlap with either VGLUT were present in every confocal image, while terminals that overlapped with VGLUT1, VGLUT2, and both VGLUTs were present in 27, 29, and 28 sections, respectively. Of the measured orexin-A putative axon terminals, 39.4 ± 4.4% overlapped with VGLUT2, 6.3 ± 2.1% overlapped with VGLUT1, and 17.7 ± 2.2% overlapped with both VGLUTs (**Figure 6d**, 1-/2+, 1+/2-, 1+/2+). The remaining 23.2 ± 6.8% of orexin-A terminals did not overlap with either VGLUT (**Figure 6d**, 1-/2-). Each of the 4 categories of orexin-A terminals were present across all OB layers with no apparent spatial distribution (**Figure 6e**).

**Figure 6:**
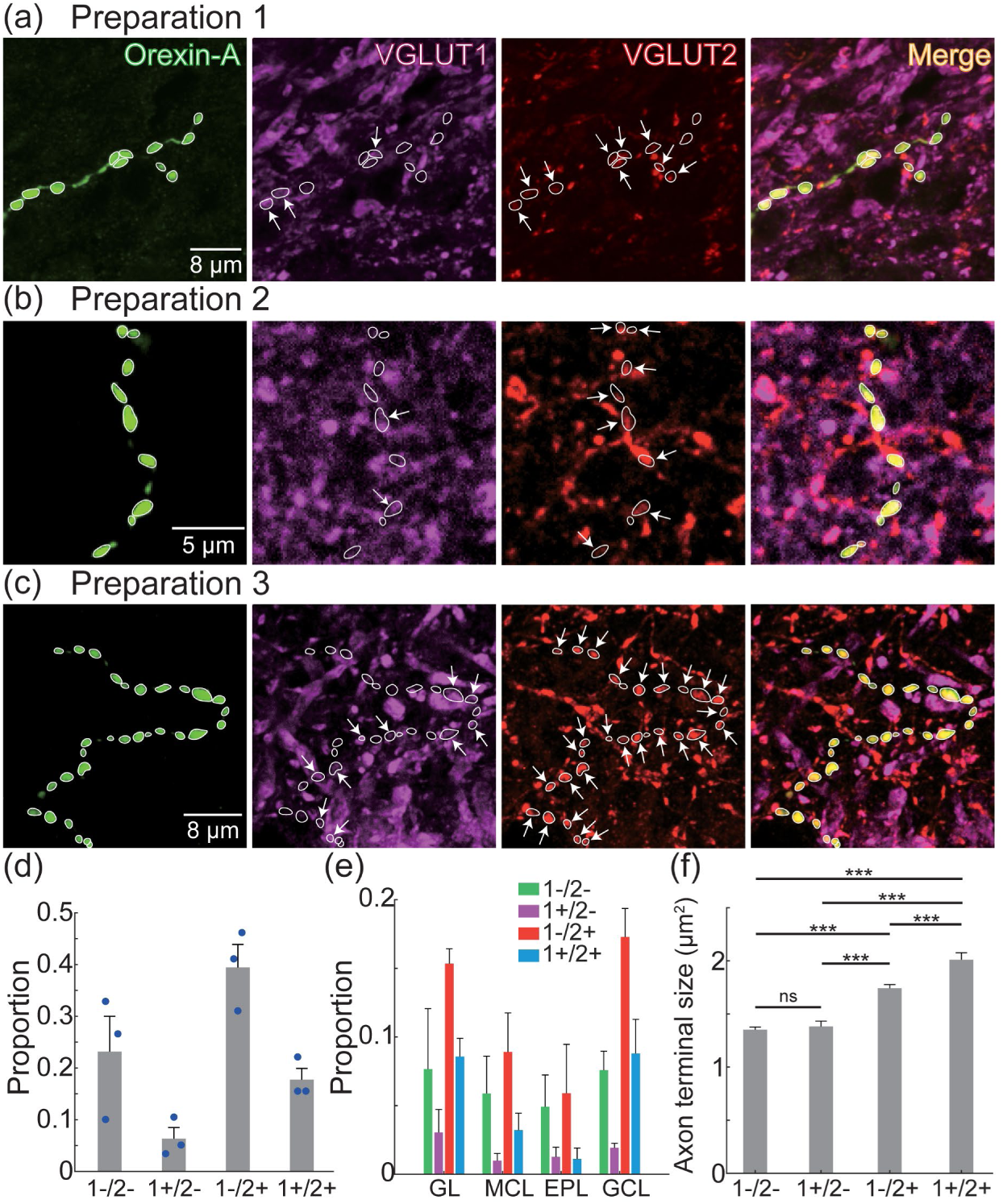
(a–c) Orexin-A, VGLUT1 (magenta), and VGLUT2 (red) expression. The images in panels a-c are z-projections of 1.2 µm, 1 µm and 2 µm, respectively. The white contours illustrate the outline of each orexin-A axon terminal. The arrows point to processes that overlap with VGLUT1 or VGLUT2. (d) The proportion of orexin-A axon terminals that did not overlap with either VGLUT (1-/2-), and that overlapped with VGLUT1 alone (1+/2-), VGLUT2 alone (1-/2+), or both VGLUTs (1+/2+). (e) Distribution of orexin-A axon terminals in different OB layers based on overlap with VGLUT1 and VGLUT2. (f) Sizes of the different orexin-A axon terminal categories.

Orexin-A terminals co-expressing VGLUT2 exhibited significantly larger areas compared to non-glutamatergic orexin-A terminals and those co-expressing only VGLUT1 (p< 0.001) (Mean ± SEM; 1-/2-: 1.35 ± 0.03 µm^2^, N = 315; 1+/2-: 1.38 ± 0.05 µm^2^, N = 87; 1-/2+: 1.74 ± 0.03 µm^2^, N = 531; 1+/2+: 2.01 ± 0.07 µm^2^, N = 238) (**Figure 6f**). Interestingly, orexin-A putative axon terminals located near each other within the same section often exhibited a mix of different overlapping properties. In some images, all orexin-A axon terminals clearly overlapped with VGLUT2 (**Figure 7a**), while other images contained both overlapping and non-overlapping processes (**Figure 7b**).

**Figure 7:**
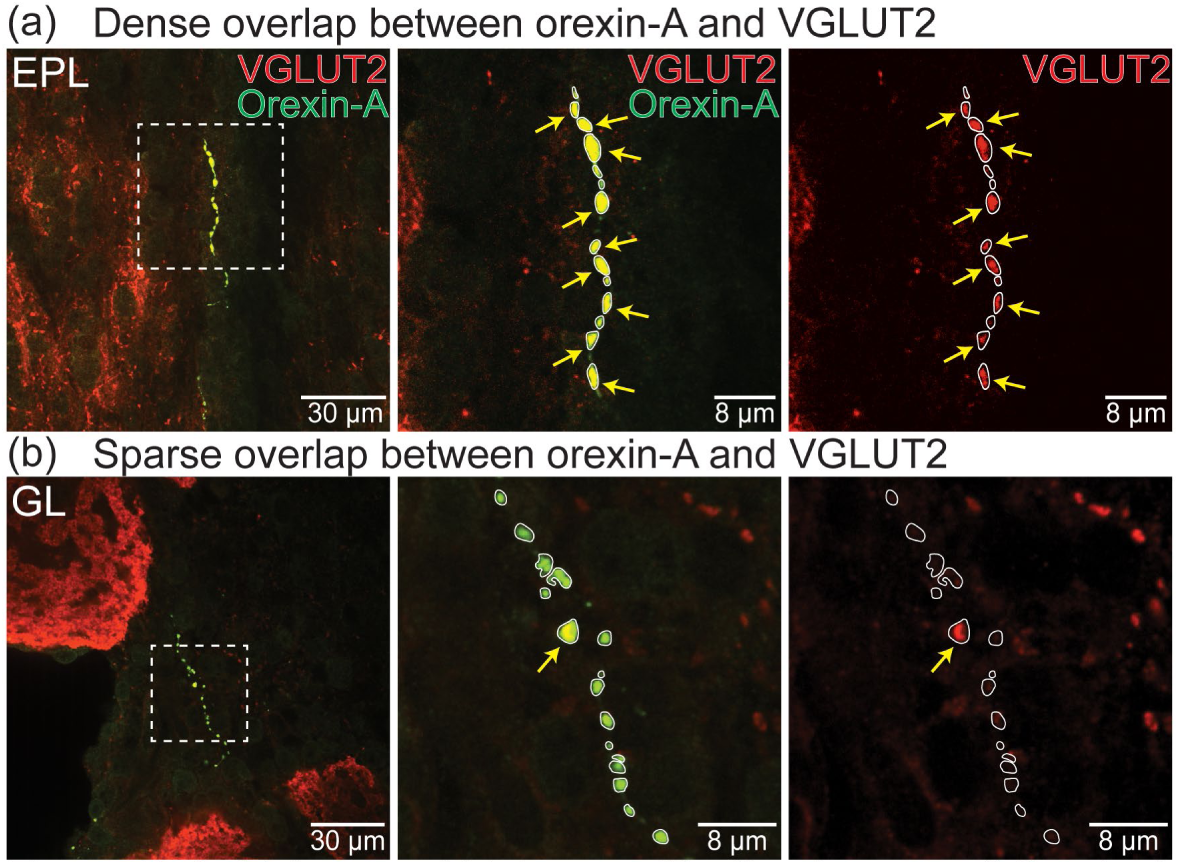
(a-b) Orexin-A (green) and VGLUT2 (red) expression in the external plexiform layer (EPL, a) and the glomerular layer (GL, b) of the OB. The arrows point to varicosities that overlap with VGLUT2.

Overlap with MCH-expressing putative axon terminals was quantified in different layers in 3 mouse preparations (N = 26 confocal images; 8-9 images per preparation, N = 661 terminals). Exemplar images illustrate the presence of overlap between MCH putative axon terminals and VGLUT2 in the OB (**Figure 8a-c**). Of the measured putative axon terminals, 72 ± 0.9% were non-glutamatergic (1-/2-), 2.6 ± 0.1% overlapped with VGLUT1 alone (1+/2-), and 25.2 ± 0.5% overlapped with VGLUT2 alone (1-/2+) (**Figure 8d**). Only 2 MCH terminals overlapped with both VGLUT isoforms and were excluded from further analysis. Each kind of labeled process was present across the different OB layers (**Figure 8e**). MCH terminals that overlapped with VGLUT2 were significantly larger than those that did not overlap with VGLUT2 (p< 0.001) and those that overlapped with VGLUT1 (Mean ± SEM; 1-/2-: 1.03 ± 0.01 µm^2^, N = 476; 1+/2-: 0.98 ± 0.06 µm^2^, N = 16; 1-/2+: 1.29 ± 0.04 µm^2^, N = 167) (**Figure 8f**).

**Figure 8:**
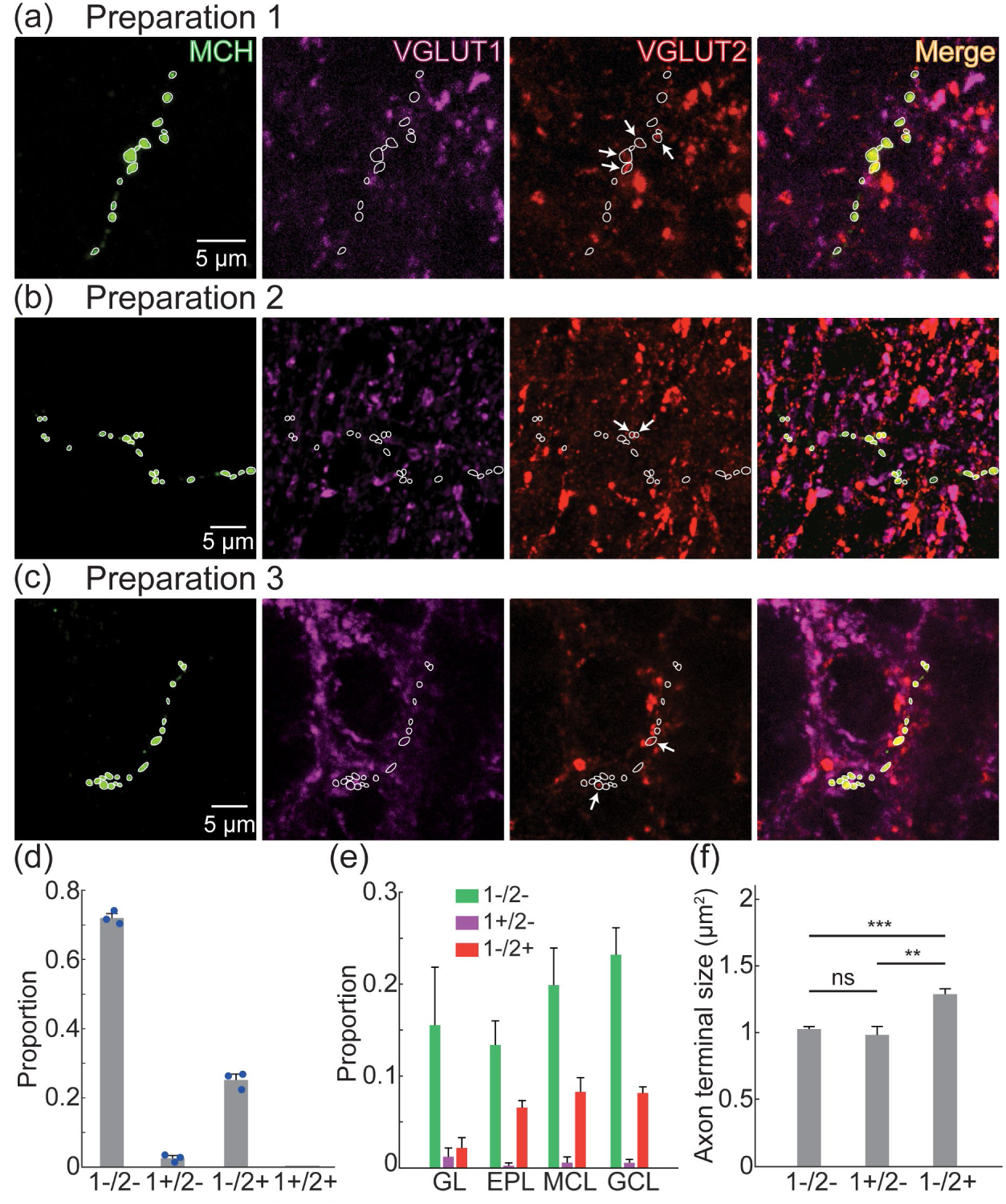
(a–c) MCH, VGLUT1 (magenta), VGLUT2 (red) expression. The images in panels a-c are z-projections of 1.2 µm, 1 µm and 2 µm, respectively. The white contours illustrate the outline of each MCH axon terminal. The arrows point to processes that overlap with VGLUT1 or VGLUT2. (d) The proportion of MCH axon terminals that did not overlap with either VGLUT (1-/2-), and that overlapped with VGLUT1 (1+/2-), VGLUT2 (1-/2+), or both VGLUTs (1+/2+). (e) Distribution of MCH axon terminals in different OB layers based on overlap with VGLUT1 and VGLUT2. (f) Sizes of MCH axon terminals.

## DISCUSSION

Here we used virally mediated tract tracing and immunohistochemistry for two different neuropeptide populations to confirm the presence of direct anterograde projections from the lateral hypothalamus to the mouse OB. We quantified the spatial location of these projections throughout the OB, finding that both orexin-A and MCH innervate all anatomical layers within the OB. Additionally, we used high-resolution confocal microscopy to demonstrate that the projections from orexin-A-and MCH-expressing neurons differently overlap with VGLUT1 and VGLUT2. The orexin-A projections to the OB are mostly glutamatergic, with 57 % overlapping with VGLUT2. In contrast, only 25 % of the MCH-expressing putative axon terminals overlapped with VGLUT2. Therefore, the OB receives a genetically heterogeneous projection from the lateral hypothalamus that includes both orexin-A and MCH-expressing neurons. Additionally, the OB input from orexin-A and MCH neurons are individually heterogeneous, with both populations expressing proteins that indicate the ability to co-release glutamate.

### Comparison with previous studies and methodological considerations

#### Anterograde projections from the orexin-expressing neurons in the lateral hypothalamus

Our results using virally-mediated tract tracing provides the first direct evidence of direct anterograde projections from orexin-A neurons in the hypothalamus to the mouse OB, confirming our previous study using a retrograde tracer (Qi et al., 2023). Although we did not quantify the spatial projection patterns of these neurons, eGFP expression was present in different layers throughout the OB (**Figure 1d-e**). Future studies are needed in which orexin-A neurons are labeled at different anterior-posterior positions within the lateral hypothalamus to investigate whether any topographic relationship exists between the hypothalamus and OB. We observed eGFP expression present in numerous brain areas (not shown), which is consistent with previous reports of orexin expression throughout the brain (Gascuel et al., 2012; Marcus et al., 2001; Nambu et al., 1999; Peyron et al., 1998; Sakurai et al., 1998). Previous studies have reported that AAV transduction using the orexin promoter can achieve a high level of specificity (González et al., 2016; Karnani et al., 2020; Y. C. Saito et al., 2013). Our study supported these results and extended them by demonstrating the effectiveness of this approach as an anterograde tracer to the OB and other brain areas (**Figure 1**).

The observation that the eGFP projections from orexin neurons did not overlap uniformly with orexin-A expression is consistent with the fact that neuropeptides are carried in secretory granules along axons of passage (**Figure 1**) (Bulgari, Zhou, Hewes, Deitcher, & Levitan, 2014; Moro, van Nifterick, Toonen, & Verhage, 2021; Yu, Liewald, Shao, Steuer Costa, & Gottschalk, 2021). In principle, immunoreactive label from both orexin-A and MCH may include both axon fibers of passage and terminals. This possibility is supported by our size measurements of all orexin-A-and MCH-expressing processes exhibiting a range of areas that are consistent with both kinds of processes (**Figure 4a, e**) (Innocenti & Caminiti, 2017; Knodel et al., 2014; Montes, Peña, DeFelipe, Herreras, & Merchan-Perez, 2015; Petrof & Sherman, 2013; Stedehouder et al., 2019). However, we cannot rule out the possibility that some of the eGFP-expressing processes in the OB that did not co-express orexin-A originated from some of the non-orexinergic eGFP-expressing neurons (**Figure 1c**, white arrows).

#### Mapping orexin-A and MCH immunoreactive processes

Previous studies have qualitatively reported the presence of orexin-A expression in the mouse OB (Gascuel et al., 2012; Qi et al., 2023). Our results extend these studies by quantitatively mapping orexin-A processes present in different OB layers across its anterior-posterior axis, finding that it is similarly distributed across each OB layer (**Figure 3c**). Additionally, we provide the first report of MCH expression in the mouse OB, finding it is non-uniformly expressed across the different OB layers.

These findings extend previous studies in the rodent OB qualitatively reporting the presence of MCH receptor mRNA, immunoreactive processes, and evidence of a direct projection (Bittencourt et al., 1992; Croizier et al., 2013; Croizier et al., 2010; Hervieu et al., 2000; Jasso et al., 2021; Y. Saito et al., 2001; Schneider et al., 2020; Skofitsch, Jacobowitz, & Zamir, 1985). Therefore, although projections from orexin-A-and MCH-expressing neurons are sparse, they are strategically positioned as a mechanism to transform sensory processing across the entire OB input-output transformation. Although the overall density of orexin-A and MCH expression was not statistically different, there was a trend for MCH to have higher levels of expression. Future retrograde tracing studies are needed to confirm the presence of direct projections from MCH-expressing neurons, and whether they provide a larger proportion of the lateral hypothalamic input to the mouse OB.

#### Overlap between neuropeptide expressing varicosities and VGLUTs

Orexin-A-and MCH-expressing neurons co-express multiple neurotransmitter systems including glutamate and dynorphin (Baldo, Daniel, Berridge, & Kelley, 2003; Blanco-Centurion et al., 2018; Chee et al., 2015; Chou et al., 2001; Mickelsen et al., 2017; Rosin et al., 2003; Yasmin et al., 2023). We provide the first report demonstrating the presence of glutamate co-expression from both neuronal populations in their axon projections to the mouse OB (**Figures 6-8**). Within these populations, we identify clear glutamatergic heterogeneity with distinct proportions of orexin-A and MCH putative axon terminals co-expressing VGLUT1 and VGLUT2 (**Figures 6** and **8**). Nearly 64% of the orexin-A varicosities overlapped with either VGLUT1, VGLUT2, or both, with the majority (∼57%) overlapping with VGLUT2. Here we also report the first measurements of the size of lateral hypothalamic varicosities innervating the OB, finding that they range between 1-1.8 µm^2^ in area.

There exist conflicting reports with respect to the degree to which orexin-A-and MCH-expressing neurons overlap with VGLUTs. Different studies have reported that between 50-90% of orexin-A-expressing neurons co-express VGLUT2, and approximately 10% co-express VGLUT1 (Blanco-Centurion et al., 2018; Mickelsen et al., 2017; Rosin et al., 2003). Therefore, our proportions are consistent with VGLUT expression within the orexin-A population. We also find that a small fraction of orexin-A projections overlap with both VGLUT1 and VGLUT2 (**Figure 6**). VGLUT1 and VGLUT2 co-expression has been reported in multiple brain areas, although no study to date has reported their co-expression in orexin-A neurons (Barroso-Chinea et al., 2007; Blanco-Centurion et al., 2018; Henny et al., 2010; Z. H. Li et al., 2020; Morimoto et al., 2003; Nakamura et al., 2007; Zander et al., 2010). There exist similarly conflicting reports with respect to co-expression of VGLUT2 in MCH neurons. There are three reports indicating that virtually all MCH neurons co-express VGLUT2, while a fourth found no overlap at all (Chee et al., 2015; Mickelsen et al., 2017; Pham et al., 2024; Schneeberger et al., 2018). Our results support the presence of VGLUT2 co-expression (**Figure 8**), yet the exact proportions remain unclear. There exist no reports of VGLUT1 co-expression in MCH-expressing neurons, which is consistent with our measurements in **Figure 8** (Mickelsen et al., 2017; Schneeberger et al., 2018). Although we are unsure of the cause of these differing results, in principle, the presence of mRNA in a cell body may not result in the protein being trafficked simultaneously to all projections from a single neuron.

#### Selection of our size threshold for quantifying VGLUT overlap

The exact proportions of overlap between neuropeptide axon terminals and VGLUT is influenced by the size threshold value we chose for orexin-A and MCH putative axon terminals (**Figure 4d**, **h**). Our results are generally consistent with a recent study showing that most overlap between orexin-A and VGLUT2 occur in large varicosities in a different brain area (Henny et al., 2010), and known sizes of glutamatergic and non-glutamatergic axon terminals in different brain areas (Bartlett, Stark, Guillery, & Smith, 2000; Covic & Sherman, 2011; Hoogland, Wouterlood, Welker, & Van der Loos, 1991; Innocenti & Caminiti, 2017; Knodel et al., 2014; J. Li, Guido, & Bickford, 2003; Montes et al., 2015; Petrof & Sherman, 2013; So, Campbell, & Lieberman, 1985). Additionally, VGLUT1 and VGLUT2 are known markers of axon terminals, and those that overlapped with orexin-A and MCH were larger than our size threshold (**Figures 4**, **6** and **8**). Although it is possible that non-glutamatergic varicosities may be systematically smaller than these, our data provide a reasonable first attempt to quantify the proportions of neuropeptide-expressing innervation to the OB that can co-release glutamate. Future studies are needed in which the lateral hypothalamic input to the OB are selectively labeled using trafficking sequences that restrict expression to axon terminals to more definitively answer this question (Beier et al., 2015; Oh et al., 2014).

#### Functional implications of neurotransmitter co-release

The observation that this pathway can likely co-release glutamate adds considerable complexity to how olfactory sensory processing can be transformed by higher brain areas (Beekly, Rupp, Burgess, & Elias, 2023). Orexin-expressing neurons that co-release glutamate do so in a spike-dependent manner where low and high spiking activity evokes glutamate and orexin release, respectively (Schöne et al., 2014; Schöne et al., 2012; Schöne, Venner, Knowles, Karnani, & Burdakov, 2011). It is unclear whether a similar mechanism exists for MCH-expressing neurons, however this provides a potential mechanism by which individual orexin-expressing neurons can modulate olfactory sensory processing on different timescales. On a broader scale, selectively knocking out VGLUT2 from MCH-expressing neurons resulted in metabolic changes that were comparable to those evoked by MCH ablation (Schneeberger et al., 2018). This suggests that glutamate signaling from neuropeptide-expressing neurons in the lateral hypothalamus play an important signaling role.

#### Conclusions

Our study provides new insights into the anatomical and neurochemical properties of hypothalamic projections to the mouse OB, highlighting their potential roles in regulating olfactory sensory processing. The presence of multiple populations of neurons in the lateral hypothalamus innervating the mouse OB provides additional support for currents models indicating that the mouse olfactory system can be modulated by neural circuits involved in state-dependent regulation at the earliest stages of sensory processing (Fadool, Tucker, & Pedarzani, 2011; Palouzier-Paulignan et al., 2012; Stark, 2024).

## Acknowledgements

This work was supported by the Karen Toffler Charitable Trust, R01 DC020519 from the National Institute of Deafness and Communication Disorders, and funding from FSU’s Dean’s Chemosensory Fellowship.

## References

Arzt, M., Deschamps, J., Schmied, C., Pietzsch, T., Schmidt, D., Tomancak, P., . . . Jug, F. (2022). LABKIT: labeling and segmentation toolkit for big image data. Frontiers in computer science, 4, 777728.

Aston-Jones, G., Smith, R. J., Sartor, G. C., Moorman, D. E., Massi, L., Tahsili-Fahadan, P., & Richardson, K. A. (2010). Lateral hypothalamic orexin/hypocretin neurons: A role in reward-seeking and addiction. Brain Res, 1314, 74–90. doi:10.1016/j.brainres.2009.09.106

Baldo, B. A., Daniel, R. A., Berridge, C. W., & Kelley, A. E. (2003). Overlapping distributions of orexin/hypocretin-and dopamine-beta-hydroxylase immunoreactive fibers in rat brain regions mediating arousal, motivation, and stress. J Comp Neurol, 464(2), 220–237. doi:10.1002/cne.10783

Barroso-Chinea, P., Castle, M., Aymerich, M. S., Pérez-Manso, M., Erro, E., Tuñon, T., & Lanciego, J. L. (2007). Expression of the mRNAs encoding for the vesicular glutamate transporters 1 and 2 in the rat thalamus. J Comp Neurol, 501(5), 703–715. doi:10.1002/cne.21265

Bartlett, E. L., Stark, J. M., Guillery, R. W., & Smith, P. H. (2000). Comparison of the fine structure of cortical and collicular terminals in the rat medial geniculate body. Neuroscience, 100(4), 811–828. doi:10.1016/s0306-4522(00)00340-7

Beekly, B. G., Rupp, A., Burgess, C. R., & Elias, C. F. (2023). Fast neurotransmitter identity of MCH neurons: Do contents depend on context? Front Neuroendocrinol, 70, 101069. doi:10.1016/j.yfrne.2023.101069

Beier, K. T., Steinberg, E. E., DeLoach, K. E., Xie, S., Miyamichi, K., Schwarz, L., . . . Luo, L. (2015). Circuit Architecture of VTA Dopamine Neurons Revealed by Systematic Input-Output Mapping. Cell, 162(3), 622–634. doi:10.1016/j.cell.2015.07.015

Berthoud, H. R., & Münzberg, H. (2011). The lateral hypothalamus as integrator of metabolic and environmental needs: from electrical self-stimulation to opto-genetics. Physiol Behav, 104(1), 29–39. doi:10.1016/j.physbeh.2011.04.051

Bittencourt, J. C., Presse, F., Arias, C., Peto, C., Vaughan, J., Nahon, J. L., . . . Sawchenko, P. E. (1992). The melanin-concentrating hormone system of the rat brain: an immuno-and hybridization histochemical characterization. J Comp Neurol, 319(2), 218–245. doi:10.1002/cne.903190204

Blanco-Centurion, C., Bendell, E., Zou, B., Sun, Y., Shiromani, P. J., & Liu, M. (2018). VGAT and VGLUT2 expression in MCH and orexin neurons in double transgenic reporter mice. IBRO Rep, 4, 44–49. doi:10.1016/j.ibror.2018.05.001

Broberger, C., De Lecea, L., Sutcliffe, J. G., & Hökfelt, T. (1998). Hypocretin/orexin-and melanin-concentrating hormone-expressing cells form distinct populations in the rodent lateral hypothalamus: relationship to the neuropeptide Y and agouti gene-related protein systems. J Comp Neurol, 402(4), 460–474.

Bulgari, D., Zhou, C., Hewes, R. S., Deitcher, D. L., & Levitan, E. S. (2014). Vesicle capture, not delivery, scales up neuropeptide storage in neuroendocrine terminals. Proc Natl Acad Sci U S A, 111(9), 3597–3601. doi:10.1073/pnas.1322170111

Caillol, M., Aïoun, J., Baly, C., Persuy, M. A., & Salesse, R. (2003). Localization of orexins and their receptors in the rat olfactory system: possible modulation of olfactory perception by a neuropeptide synthetized centrally or locally. Brain Res, 960(1-2), 48–61. doi:10.1016/s0006-8993(02)03755-1

Chee, M. J., Arrigoni, E., & Maratos-Flier, E. (2015). Melanin-concentrating hormone neurons release glutamate for feedforward inhibition of the lateral septum. J Neurosci, 35(8), 3644–3651. doi:10.1523/JNEUROSCI.4187-14.2015

Chelette, B. M., Loeven, A. M., Gatlin, D. N., Landi Conde, D. R., Huffstetler, C. M., Qi, M., & Fadool, D. A. (2021). Consumption of dietary fat causes loss of olfactory sensory neurons and associated circuitry that is not mitigated by voluntary exercise in mice. J Physiol. doi:10.1113/JP282112

Chou, T. C., Lee, C. E., Lu, J., Elmquist, J. K., Hara, J., Willie, J. T., . . . Scammell, T. E. (2001). Orexin (hypocretin) neurons contain dynorphin. J Neurosci, 21(19), RC168.

Concetti, C., Peleg-Raibstein, D., & Burdakov, D. (2024). Hypothalamic MCH Neurons: From Feeding to Cognitive Control. Function (Oxf), 5(1), zqad059. doi:10.1093/function/zqad059

Covic, E. N., & Sherman, S. M. (2011). Synaptic properties of connections between the primary and secondary auditory cortices in mice. Cereb Cortex, 21(11), 2425–2441. doi:10.1093/cercor/bhr029

Croizier, S., Cardot, J., Brischoux, F., Fellmann, D., Griffond, B., & Risold, P. Y. (2013). The vertebrate diencephalic MCH system: a versatile neuronal population in an evolving brain. Front Neuroendocrinol, 34(2), 65–87. doi:10.1016/j.yfrne.2012.10.001

Croizier, S., Franchi-Bernard, G., Colard, C., Poncet, F., La Roche, A., & Risold, P. Y. (2010). A comparative analysis shows morphofunctional differences between the rat and mouse melanin-concentrating hormone systems. PLoS One, 5(11), e15471. doi:10.1371/journal.pone.0015471

Dawson, M., Terstege, D. J., Jamani, N., Tsutsui, M., Pavlov, D., Bugescu, R., . . . Sargin, D. (2023). Hypocretin/orexin neurons encode social discrimination and exhibit a sex-dependent necessity for social interaction. Cell Rep, 42(7), 112815. doi:10.1016/j.celrep.2023.112815

Diniz, G. B., & Bittencourt, J. C. (2017). The Melanin-Concentrating Hormone as an Integrative Peptide Driving Motivated Behaviors. Front Syst Neurosci, 11, 32. doi:10.3389/fnsys.2017.00032

El Mestikawy, S., Wallén-Mackenzie, A., Fortin, G. M., Descarries, L., & Trudeau, L. E. (2011). From glutamate co-release to vesicular synergy: vesicular glutamate transporters. Nat Rev Neurosci, 12(4), 204–216. doi:10.1038/nrn2969

Fadool, D. A., Tucker, K., & Pedarzani, P. (2011). Mitral cells of the olfactory bulb perform metabolic sensing and are disrupted by obesity at the level of the Kv1.3 ion channel. PLoS One, 6(9), e24921. doi:10.1371/journal.pone.0024921

Fardone, E., Celen, A. B., Schreiter, N. A., Thiebaud, N., Cooper, M. L., & Fadool, D. A. (2019). Loss of odor-induced c-Fos expression of juxtaglomerular activity following maintenance of mice on fatty diets. J Bioenerg Biomembr, 51(1), 3–13. doi:10.1007/s10863-018-9769-5

Franklin, K., & Paxinos, G. (2008). The Mouse Brain in Stereotaxic Coordinates, Compact - 3rd Edition.

Fremeau, R. T., Troyer, M. D., Pahner, I., Nygaard, G. O., Tran, C. H., Reimer, R. J., . . . Edwards, R. H. (2001). The expression of vesicular glutamate transporters defines two classes of excitatory synapse. Neuron, 31(2), 247–260. doi:10.1016/s0896-6273(01)00344-0

Fu, O., Iwai, Y., Narukawa, M., Ishikawa, A. W., Ishii, K. K., Murata, K., . . . Nakajima, K. I. (2019). Hypothalamic neuronal circuits regulating hunger-induced taste modification. Nat Commun, 10(1), 4560. doi:10.1038/s41467-019-12478-x

Gabellec, M. M., Panzanelli, P., Sassoè-Pognetto, M., & Lledo, P. M. (2007). Synapse-specific localization of vesicular glutamate transporters in the rat olfactory bulb. Eur J Neurosci, 25(5), 1373–1383. doi:10.1111/j.1460-9568.2007.05400.x

Gascuel, J., Lemoine, A., Rigault, C., Datiche, F., Benani, A., Penicaud, L., & Lopez-Mascaraque, L. (2012). Hypothalamus-olfactory system crosstalk: orexin a immunostaining in mice. Front Neuroanat, 6, 44. doi:10.3389/fnana.2012.00044

González, J. A., Jensen, L. T., Iordanidou, P., Strom, M., Fugger, L., & Burdakov, D. (2016). Inhibitory Interplay between Orexin Neurons and Eating. Curr Biol, 26(18), 2486–2491. doi:10.1016/j.cub.2016.07.013

Gross-Isseroff, R., & Lancet, D. (1988). Concentration-dependent changes of perceived odor quality. Chemical Senses, 13(2), 191–204. doi:10.1093/chemse/13.2.191

Hardy, A. B., Aïoun, J., Baly, C., Julliard, K. A., Caillol, M., Salesse, R., & Duchamp-Viret, P. (2005). Orexin A modulates mitral cell activity in the rat olfactory bulb: patch-clamp study on slices and immunocytochemical localization of orexin receptors. Endocrinology, 146(9), 4042–4053. doi:10.1210/en.2005-0020

Henny, P., Brischoux, F., Mainville, L., Stroh, T., & Jones, B. E. (2010). Immunohistochemical evidence for synaptic release of glutamate from orexin terminals in the locus coeruleus. Neuroscience, 169(3), 1150–1157. doi:10.1016/j.neuroscience.2010.06.003

Hervieu, G. J., Cluderay, J. E., Harrison, D., Meakin, J., Maycox, P., Nasir, S., & Leslie, R. A. (2000). The distribution of the mRNA and protein products of the melanin-concentrating hormone (MCH) receptor gene, slc-1, in the central nervous system of the rat. Eur J Neurosci, 12(4), 1194–1216. doi:10.1046/j.1460-9568.2000.00008.x

Herzog, E., Bellenchi, G. C., Gras, C., Bernard, V., Ravassard, P., Bedet, C., . . . El Mestikawy, S. (2001). The existence of a second vesicular glutamate transporter specifies subpopulations of glutamatergic neurons. J Neurosci, 21(22), RC181. doi:10.1523/JNEUROSCI.21-22-j0001.2001

Homma, R., Cohen, L. B., Kosmidis, E. K., & Youngentob, S. L. (2009). Perceptual stability during dramatic changes in olfactory bulb activation maps and dramatic declines in activation amplitudes. Eur J Neurosci, 29(5), 1027–1034. doi:10.1111/j.1460-9568.2009.06644.x

Hoogland, P. V., Wouterlood, F. G., Welker, E., & Van der Loos, H. (1991). Ultrastructure of giant and small thalamic terminals of cortical origin: a study of the projections from the barrel cortex in mice using Phaseolus vulgaris leuco-agglutinin (PHA-L). Exp Brain Res, 87(1), 159–172. doi:10.1007/BF00228517

In’t Zandt, E. E., Cansler, H. L., Denson, H. B., & Wesson, D. W. (2019). Centrifugal Innervation of the Olfactory Bulb: A Reappraisal. eNeuro, 6(1). doi:10.1523/eneuro.0390-18.2019

Innocenti, G. M., & Caminiti, R. (2017). Axon diameter relates to synaptic bouton size: structural properties define computationally different types of cortical connections in primates. Brain Struct Funct, 222(3), 1169–1177. doi:10.1007/s00429-016-1266-1

Jasso, K. R., Kamba, T. K., Zimmerman, A. D., Bansal, R., Engle, S. E., Everett, T., . . . McIntyre, J. C. (2021). An N-terminal fusion allele to study melanin concentrating hormone receptor 1. Genesis, 59(7-8), e23438. doi:10.1002/dvg.23438

Julliard, A. K., Al Koborssy, D., Fadool, D. A., & Palouzier-Paulignan, B. (2017). Nutrient Sensing: Another Chemosensitivity of the Olfactory System. Front Physiol, 8, 468. doi:10.3389/fphys.2017.00468

Karnani, M. M., Schöne, C., Bracey, E. F., González, J. A., Viskaitis, P., Li, H. T., . . . Burdakov, D. (2020). Role of spontaneous and sensory orexin network dynamics in rapid locomotion initiation. Prog Neurobiol, 187, 101771. doi:10.1016/j.pneurobio.2020.101771

Knodel, M. M., Geiger, R., Ge, L., Bucher, D., Grillo, A., Wittum, G., . . . Queisser, G. (2014). Synaptic bouton properties are tuned to best fit the prevailing firing pattern. Front Comput Neurosci, 8, 101. doi:10.3389/fncom.2014.00101

Kolling, L. J., Tatti, R., Lowry, T., Loeven, A. M., Fadool, J. M., & Fadool, D. A. (2022). Modulating the Excitability of Olfactory Output Neurons Affects Whole-Body Metabolism. J Neurosci, 42(30), 5966–5990. doi:10.1523/JNEUROSCI.0190-22.2022

Li, J., Guido, W., & Bickford, M. E. (2003). Two distinct types of corticothalamic EPSPs and their contribution to short-term synaptic plasticity. J Neurophysiol, 90(5), 3429–3440. doi:10.1152/jn.00456.2003

Li, Z. H., Zhang, C. K., Qiao, Y., Ge, S. N., Zhang, T., & Li, J. L. (2020). Coexpression of VGLUT1 and VGLUT2 in precerebellar neurons in the lateral reticular nucleus of the rat. Brain Res Bull, 162, 94–106. doi:10.1016/j.brainresbull.2020.06.008

Marcus, J. N., Aschkenasi, C. J., Lee, C. E., Chemelli, R. M., Saper, C. B., Yanagisawa, M., & Elmquist, J. K. (2001). Differential expression of orexin receptors 1 and 2 in the rat brain. J Comp Neurol, 435(1), 6–25. doi:10.1002/cne.1190

Mickelsen, L. E., Kolling, F. W., Chimileski, B. R., Fujita, A., Norris, C., Chen, K., . . . Jackson, A. C. (2017). Neurochemical Heterogeneity Among Lateral Hypothalamic Hypocretin/Orexin and Melanin-Concentrating Hormone Neurons Identified Through Single-Cell Gene Expression Analysis. eNeuro, 4(5). doi:10.1523/ENEURO.0013-17.2017

Montes, J., Peña, J. M., DeFelipe, J., Herreras, O., & Merchan-Perez, A. (2015). The influence of synaptic size on AMPA receptor activation: a Monte Carlo model. PLoS One, 10(6), e0130924. doi:10.1371/journal.pone.0130924

Morimoto, R., Hayashi, M., Yatsushiro, S., Otsuka, M., Yamamoto, A., & Moriyama, Y. (2003). Co-expression of vesicular glutamate transporters (VGLUT1 and VGLUT2) and their association with synaptic-like microvesicles in rat pinealocytes. J Neurochem, 84(2), 382–391. doi:10.1046/j.1471-4159.2003.01532.x

Moro, A., van Nifterick, A., Toonen, R. F., & Verhage, M. (2021). Dynamin controls neuropeptide secretion by organizing dense-core vesicle fusion sites. Sci Adv, 7(21). doi:10.1126/sciadv.abf0659

Nakamura, K., Hioki, H., Fujiyama, F., & Kaneko, T. (2005). Postnatal changes of vesicular glutamate transporter (VGluT)1 and VGluT2 immunoreactivities and their colocalization in the mouse forebrain. J Comp Neurol, 492(3), 263–288. doi:10.1002/cne.20705

Nakamura, K., Watakabe, A., Hioki, H., Fujiyama, F., Tanaka, Y., Yamamori, T., & Kaneko, T. (2007). Transiently increased colocalization of vesicular glutamate transporters 1 and 2 at single axon terminals during postnatal development of mouse neocortex: a quantitative analysis with correlation coefficient. Eur J Neurosci, 26(11), 3054–3067. doi:10.1111/j.1460-9568.2007.05868.x

Nambu, T., Sakurai, T., Mizukami, K., Hosoya, Y., Yanagisawa, M., & Goto, K. (1999). Distribution of orexin neurons in the adult rat brain. Brain Res, 827(1-2), 243–260. doi:10.1016/s0006-8993(99)01336-0

Oh, S. W., Harris, J. A., Ng, L., Winslow, B., Cain, N., Mihalas, S., . . . Zeng, H. (2014). A mesoscale connectome of the mouse brain. Nature, 508(7495), 207–214. doi:10.1038/nature13186

Ohmomo, H., Ina, A., Yoshida, S., Shutoh, F., Ueda, S., & Hisano, S. (2009). Postnatal changes in expression of vesicular glutamate transporters in the main olfactory bulb of the rat. Neuroscience, 160(2), 419–426. doi:10.1016/j.neuroscience.2009.02.048

Palouzier-Paulignan, B., Lacroix, M. C., Aime, P., Baly, C., Caillol, M., Congar, P., . . . Fadool, D. A. (2012). Olfaction under metabolic influences. Chem Senses, 37(9), 769–797. doi:10.1093/chemse/bjs059

Petrof, I., & Sherman, S. M. (2013). Functional significance of synaptic terminal size in glutamatergic sensory pathways in thalamus and cortex. J Physiol, 591(13), 3125–3131. doi:10.1113/jphysiol.2012.247619

Peyron, C., Faraco, J., Rogers, W., Ripley, B., Overeem, S., Charnay, Y., . . . Mignot, E. (2000). A mutation in a case of early onset narcolepsy and a generalized absence of hypocretin peptides in human narcoleptic brains. Nat Med, 6(9), 991–997. doi:10.1038/79690

Peyron, C., Tighe, D. K., van den Pol, A. N., de Lecea, L., Heller, H. C., Sutcliffe, J. G., & Kilduff, T. S. (1998). Neurons containing hypocretin (orexin) project to multiple neuronal systems. J Neurosci, 18(23), 9996–10015.

Pham, X. T., Abe, Y., Mukai, Y., Ono, D., Tanaka, K. F., Ohmura, Y., . . . Yamanaka, A. (2024). Glutamatergic signaling from melanin-concentrating hormone-producing neurons: A requirement for memory regulation, but not for metabolism control. PNAS Nexus, 3(7), pgae275. doi:10.1093/pnasnexus/pgae275

Qi, M., Fadool, D. A., & Storace, D. A. (2023). An anatomically distinct subpopulation of orexin neurons project from the lateral hypothalamus to the olfactory bulb. J Comp Neurol. doi:10.1002/cne.25518

Qu, D., Ludwig, D. S., Gammeltoft, S., Piper, M., Pelleymounter, M. A., Cullen, M. J., . . . Maratos-Flier, E. (1996). A role for melanin-concentrating hormone in the central regulation of feeding behaviour. Nature, 380(6571), 243–247. doi:10.1038/380243a0

Rao, Y., Lu, M., Ge, F., Marsh, D. J., Qian, S., Wang, A. H., . . . Gao, X. B. (2008). Regulation of synaptic efficacy in hypocretin/orexin-containing neurons by melanin concentrating hormone in the lateral hypothalamus. J Neurosci, 28(37), 9101–9110. doi:10.1523/JNEUROSCI.1766-08.2008

Rokni, D., Hemmelder, V., Kapoor, V., & Murthy, V. N. (2014). An olfactory cocktail party: figure-ground segregation of odorants in rodents. Nat Neurosci, 17(9), 1225–1232. doi:10.1038/nn.3775

Rosin, D. L., Weston, M. C., Sevigny, C. P., Stornetta, R. L., & Guyenet, P. G. (2003). Hypothalamic orexin (hypocretin) neurons express vesicular glutamate transporters VGLUT1 or VGLUT2. J Comp Neurol, 465(4), 593–603. doi:10.1002/cne.10860

Saito, Y., Cheng, M., Leslie, F. M., & Civelli, O. (2001). Expression of the melanin-concentrating hormone (MCH) receptor mRNA in the rat brain. J Comp Neurol, 435(1), 26–40. doi:10.1002/cne.1191

Saito, Y., & Nagasaki, H. (2008). The melanin-concentrating hormone system and its physiological functions. Results Probl Cell Differ, 46, 159–179. doi:10.1007/400_2007_052

Saito, Y. C., Tsujino, N., Hasegawa, E., Akashi, K., Abe, M., Mieda, M., . . . Sakurai, T. (2013). GABAergic neurons in the preoptic area send direct inhibitory projections to orexin neurons. Front Neural Circuits, 7, 192. doi:10.3389/fncir.2013.00192

Sakurai, T., Amemiya, A., Ishii, M., Matsuzaki, I., Chemelli, R. M., Tanaka, H., . . . Yanagisawa, M. (1998). Orexins and orexin receptors: a family of hypothalamic neuropeptides and G protein-coupled receptors that regulate feeding behavior. Cell, 92(5), 1 page following 696. doi:10.1016/s0092-8674(02)09256-5

Schneeberger, M., Tan, K., Nectow, A. R., Parolari, L., Caglar, C., Azevedo, E., . . . Friedman, J. M. (2018). Functional analysis reveals differential effects of glutamate and MCH neuropeptide in MCH neurons. Mol Metab, 13, 83–89. doi:10.1016/j.molmet.2018.05.001

Schneider, N. Y., Chaudy, S., Epstein, A. L., Viollet, C., Benani, A., Penicaud, L., . . . Gascuel, J. (2020). Centrifugal projections to the main olfactory bulb revealed by transsynaptic retrograde tracing in mice. J Comp Neurol, 528(11), 1805–1819. doi:10.1002/cne.24846

Schöne, C., Apergis-Schoute, J., Sakurai, T., Adamantidis, A., & Burdakov, D. (2014). Coreleased orexin and glutamate evoke nonredundant spike outputs and computations in histamine neurons. Cell Rep, 7(3), 697–704. doi:10.1016/j.celrep.2014.03.055

Schöne, C., Cao, Z. F., Apergis-Schoute, J., Adamantidis, A., Sakurai, T., & Burdakov, D. (2012). Optogenetic probing of fast glutamatergic transmission from hypocretin/orexin to histamine neurons in situ. J Neurosci, 32(36), 12437–12443. doi:10.1523/JNEUROSCI.0706-12.2012

Schöne, C., Venner, A., Knowles, D., Karnani, M. M., & Burdakov, D. (2011). Dichotomous cellular properties of mouse orexin/hypocretin neurons. J Physiol, 589(Pt 11), 2767–2779. doi:10.1113/jphysiol.2011.208637

Shibata, M., Mondal, M. S., Date, Y., Nakazato, M., Suzuki, H., & Ueta, Y. (2008). Distribution of orexins-containing fibers and contents of orexins in the rat olfactory bulb. Neurosci Res, 61(1), 99–105. doi:10.1016/j.neures.2008.01.017

Shipley, M. T., & Adamek, G. D. (1984). The connections of the mouse olfactory bulb: a study using orthograde and retrograde transport of wheat germ agglutinin conjugated to horseradish peroxidase. Brain Res Bull, 12(6), 669–688.

Skofitsch, G., Jacobowitz, D. M., & Zamir, N. (1985). Immunohistochemical localization of a melanin concentrating hormone-like peptide in the rat brain. Brain Res Bull, 15(6), 635–649. doi:10.1016/0361-9230(85)90213-8

So, K. F., Campbell, G., & Lieberman, A. R. (1985). Synaptic organization of the dorsal lateral geniculate nucleus in the adult hamster. An electron microscope study using degeneration and horseradish peroxidase tracing techniques. Anat Embryol (Berl), 171(2), 223–234. doi:10.1007/BF00341417

Stark, R. (2024). The olfactory bulb: A neuroendocrine spotlight on feeding and metabolism. J Neuroendocrinol, 36(6), e13382. doi:10.1111/jne.13382

Stedehouder, J., Brizee, D., Slotman, J. A., Pascual-Garcia, M., Leyrer, M. L., Bouwen, B. L., . . . Kushner, S. A. (2019). Local axonal morphology guides the topography of interneuron myelination in mouse and human neocortex. Elife, 8. doi:10.7554/eLife.48615

Subramanian, K. S., Lauer, L. T., Hayes, A. M. R., Décarie-Spain, L., McBurnett, K., Nourbash, A. C., . . . Kanoski, S. E. (2023). Hypothalamic melanin-concentrating hormone neurons integrate food-motivated appetitive and consummatory processes in rats. Nat Commun, 14(1), 1755. doi:10.1038/s41467-023-37344-9

Takamori, S., Rhee, J. S., Rosenmund, C., & Jahn, R. (2000). Identification of a vesicular glutamate transporter that defines a glutamatergic phenotype in neurons. Nature, 407(6801), 189–194. doi:10.1038/35025070

Thannickal, T. C., Moore, R. Y., Nienhuis, R., Ramanathan, L., Gulyani, S., Aldrich, M., . . . Siegel, J. M. (2000). Reduced number of hypocretin neurons in human narcolepsy. Neuron, 27(3), 469–474. doi:10.1016/s0896-6273(00)00058-1

Timper, K., & Bruning, J. C. (2017). Hypothalamic circuits regulating appetite and energy homeostasis: pathways to obesity. Dis Model Mech, 10(6), 679–689. doi:10.1242/dmm.026609

Uchida, N., & Mainen, Z. F. (2007). Odor concentration invariance by chemical ratio coding. Front Syst Neurosci, 1, 3. doi:10.3389/neuro.06.003.2007

Wheeler, D. S., Wan, S., Miller, A., Angeli, N., Adileh, B., Hu, W., & Holland, P. C. (2014). Role of lateral hypothalamus in two aspects of attention in associative learning. Eur J Neurosci, 40(2), 2359–2377. doi:10.1111/ejn.12592

Yamanaka, A., Beuckmann, C. T., Willie, J. T., Hara, J., Tsujino, N., Mieda, M., . . . Sakurai, T. (2003). Hypothalamic orexin neurons regulate arousal according to energy balance in mice. Neuron, 38(5), 701–713. doi:10.1016/s0896-6273(03)00331-3

Yasmin, N., Collier, A. D., Karatayev, O., Abdulai, A. R., Yu, B., Fam, M., . . . Leibowitz, S. F. (2023). Subpopulations of hypocretin/orexin neurons differ in measures of their cell proliferation, dynorphin co-expression, projections, and response to embryonic ethanol exposure. Sci Rep, 13(1), 8448. doi:10.1038/s41598-023-35432-w

Yu, S. C., Liewald, J. F., Shao, J., Steuer Costa, W., & Gottschalk, A. (2021). Synapsin Is Required for Dense Core Vesicle Capture and cAMP-Dependent Neuropeptide Release. J Neurosci, 41(19), 4187–4201. doi:10.1523/JNEUROSCI.2631-20.2021

Zander, J. F., Münster-Wandowski, A., Brunk, I., Pahner, I., Gómez-Lira, G., Heinemann, U., . . . Ahnert-Hilger, G. (2010). Synaptic and vesicular coexistence of VGLUT and VGAT in selected excitatory and inhibitory synapses. J Neurosci, 30(22), 7634–7645. doi:10.1523/JNEUROSCI.0141-10.2010

